# Comparative genomic analysis of a Colombian strain of *Leptospira santarosai* serogroup Autumnalis serovar Alice and inference of its virulence factors

**DOI:** 10.1101/2024.08.09.607408

**Authors:** Rafael Guillermo Villarreal-Julio, Brayan Ordoñez, Jonny Andrés Yepes-Blandón

## Abstract

**Introduction:** The study aimed to characterize the genome of a Colombian strain of *Leptospira santarosai* to infer bacterial virulence factors.

**Materials and methods:** Under the approach of a quantitative research, an isolate of Colombian origin of *L. santarosai* was sequenced using new generation 454 GS FLX Titanium sequencers. Subsequently, bioinformatics programs were used for the physical description of the genome (number of genes, size), open reading frame prediction, prediction of proteins and their orthologs in other pathogenic species, of intermediate pathogenicity and non-pathogenic, cellular localization of proteins.

**Results:** The assembly of the spirochete genome was achieved *Leptospira santarosai* serogroup Autumnalis serovar Aliceand the description of the genetic, structural and functional characteristics of its genes. It is concluded that the core-genome of the isolate is composed of 1747 proteins, which are common to all *L. santarosai* strains available in GenBank. It was determined that it has a total of 4,138 proteins, 141 of which are unique to its genome. The possible role of the virulence factors found in the Colombian isolate was identified and described.

**Conclusions:** The study contributes to the understanding of the pathophysiological mechanisms induced by *Leptospira*.

## Introduction

Leptospirosis is a re-emerging zoonosis with worldwide distribution.^1.2^ caused by spirochetes of the genus *Leptospira*. The World Health Organization (WHO) estimated a fatality rate ranging from 5 to 30%, and an incidence rate of 4 to 100 cases per one hundred thousand inhabitants in tropical and subtropical countries.^3^. In Colombia, through SIVIGILA in epidemiological week 12 of 2016, 645 suspected cases of leptospirosis were reported^4^. These reports demonstrate the current circulation of *Leptospira* spp in this territory, however, the real situation of the disease is unknown in the vast majority of regions of the country.

The symptoms of leptospirosis are varied and include fever, jaundice, kidney failure and/or pulmonary hemorrhage, culminating in multiple organ failure^5^. Pathogenic *Leptospira* species are transmitted to humans by direct contact with reservoirs or through water contaminated with urine from infected animals^6^.

Kidney injury is the early manifestation of leptospirosis and occurs a few days after infection, presenting tubulointerstitial nephritis that can trigger damage to the renal blood vessels and the structure of the kidney, while in chronic infection tubulointerstitial nephritis occurs, which progresses to fibrosis and eventually leads to kidney failure^7.8^.

In chronic infection, the bacteria harbors in the renal tubules and the mechanisms by which it induces renal pathogenesis are not completely clarified and the virulence mechanisms that contribute to this pathogenicity are not completely identified, although various factors such as lipopolysaccharides (LPS), hemolysins, adhesion proteins, endotoxins among others have been proposed^9,10^.

The genus *Leptospira* has a wide genetic diversity that includes groups of species that comprise 22 species between pathogenic, intermediate and saprophytic.^11^.

All *Leptospira* strains generally present two circular chromosomes, one called the major chromosome or CI and the other minor chromosome or CII, and all the genomes present notable differences in terms of size, genomic composition, number of genes and expressed proteins, with evidence of large-scale genomic rearrangements^12–14^.

The term virulence factor is used to describe proteins, structures (LPS, carbohydrates) or phenotypes (motility) that are essential to produce the disease, or that have been shown to enhance the disease by interacting with host proteins. Various virulence factors have been identified in pathogenic *Leptospira* species including lipopolysaccharides, lipoproteins, outer membrane proteins (OMPs) and wall components which are functionally and structurally important for nutrient acquisition, signaling, environmental adaptation, survival and immunogenicity.^15,16^. Other virulence factors consist of those that allow it to fixate in the renal tubules and generate colonization or renal persistence in reservoirs. Some of them intervene in bacterial motility and chemotaxis so that the pathogen reaches and remains in its target organ to subsequently remain latent and causing infection.^17^.

Within the pathogenic species of *Leptospira* we find *L. santarosai* serovar Shermani, a strain isolated in Taiwan, whose genome has a size of 3,983,611 bp, base pairs, composed of two circular chromosomes, one larger than 3,659,905 bp and one smaller than 323 706 pb18. The genome of *L. santarosai* serovar Shermani has a GC content of 41.82%. It presents 4033 computationally predicted genes, it has a percentage of coding DNA of 92.5%, these genes have a function of motility, chemotaxis, and formation of lipoproteins and outer membrane proteins mainly, although genes have also been found that are also *L. interrogans, L. borgpetersenii and L. biflexa* for replication, recombination, repair, cell cycle control and cell division^5.18^. Its genome also has 48 transposases, which favor genetic diversity and adaptation to different environmental conditions, which is evidence of genetic alterations and reorganization during its evolutionary history. It has 5 ribosomal RNA (rRNA) genes and 37 transfer RNA genes^5.18^. *L. santarosai* serovar Shermani is evolutionarily most related to *L. borgpetersenii*. The analysis of functional groups reveals that in *L.* santorosai serovar Shermani between 1.9% and 9.4% of the genes sequence for cell wall, membrane, biogenesis of defense mechanisms, and in *L. interrogans* this comparison resulted for the same genes a lower percentage ranging from 1.5% to 8%^18^.

Although we currently know the importance of knowledge at the genetic level, there is great ignorance about the pathogenesis of *Leptospira* from Colombia, to such an extent that there is not a single complete genome sequence of isolates present and reported in Colombia, such as Knowledge would be directly framed in its biology, genetic and structural mechanisms, which can be elucidated by knowing its genome.

In this study, the genome of a Colombian strain of *Leptospira santarosai* serogroup Autumnalis serovar Alice, a species never before reported in Colombia, has been sequenced. This strain was collected from a previous study carried out in the Urabá region of Antioquia and was isolated from a male patient. 13 years old, from Apartadó4. This strain is virulent and presents varied and non-specific symptoms in the host, and also shows less severity in terms of virulence than other pathogenic species.

In the Colombian strain *L. santarosai*, virulence factors were identified through a comparative genomic analysis between the different groups of pathogenic, intermediate and saprophytic *Leptospira*.

The new characteristics of *Leptospira santarosai* serogroup Autumnalis serovar Alice found in this study may contribute to the understanding of the pathophysiological mechanisms of *Leptospira*. Comparative genomic characteristics, especially between human pathogenic species, will allow us to identify the virulence factors that the bacteria need to infect, persist and cause the disease.

## Materials and methods

### Leptospira strain

The strain *L. santarosai* serogroup Autumnalis serovar Alice of Colombian origin was isolated from a male patient who was 13 years old, and was a native of Apartadó, Colombia, who had been suffering from general malaise for three days, who in the Clinical inspection revealed renal symptoms associated with the clinical presentation called Weil syndrome. The main symptomatology was a febrile condition that consisted of fever, headache, ocular itching with conjunctival injection, “stoned” tarsal conjunctiva, “periocular wheals” and persistent cough, but without meningeal signs^4^.

### Diagnosis

A rapid serological test was performed to detect IgM for *Leptospira*. The serological test gave a positive result and the patient was ordered doxycycline in a dose of 200 mg per day over a period of 14 days, achieving the elimination of symptoms on the fourth day of treatment provided.

### Blood culture

A blood culture was performed in Fletcher’s semisolid medium. This was positive, which confirmed the diagnosis of leptospirosis in the patient, and this culture was used to extract genetic material to be used in bacterial genotyping and to obtain subcultures using the methods frequently used in biological assays.

### Isolation

To carry out subcultures, the isolates were preserved in picks, which were carried out every two weeks, over a period of around 6 to 8 months on average. Bacterial cultures were performed in EMJH liquid medium (Ellinghausen-McCullough-Johnson-Harris, BectonDickinson-Biosciences) supplemented with 10% commercial enrichment medium (Becton-Dickinson-Biosciences) and were incubated between 26°C and 30°C. C under aerobic conditions.

### Genome sequencing, assembly and annotation

DNA was extracted from the culture of the *L. santarosai* strain isolated from the patient using the kit: DNeasy Qiagen, following the manufacturer’s instructions. It was subsequently subjected to sequencing using dual next generation sequencing platforms (NGS: next-generation sequencing) 454 GS FLX Titanium (Roche, Branford, USA) and Illumina HiSeq 2000 (Illumina Inc., San Diego, CA, USA), thus obtaining better coverage of the genome sequence. An RS Pacific Biosciences sequencer (PacBio; Pacific Biosciences, Menlo Park, USA) was used. De novo assembly was performed using Celera Assembler v. 7.0. The order and orientation of these contigs were corroborated using an optical mapping system (Opgen Technologies Inc., Madison, WI, USA).

This strain is part of the *Leptospira* and Human Health Genomics Project, as part of the Infectious Diseases Genome Sequencing Center. DNA and/or bacterial genomic material are available in the BEI resource repository (http://beiresources.org). This sequencing project has been funded with federal funds from the National Institute of Allergy and Infectious Diseases of the National Institutes of Health, Department of Health and Human Services through the Genomic Sequencing Centers for Infectious Diseases under contract number HHSN272200900007C. The sequence was submitted by the J. Craig Venter Institute and the annotation of the contigs was updated in March 2013.

### Nucleotide sequence accession number

The genome sequence of the Colombian strain *L. santarosai* serogroup Autumnalis serovar Alice has been deposited in DDBJ/EMBL/GenBank with the accession number AKWS00000000.2.

### Comparative genomic analysis

The circular genome map was generated from the Viewer server (CGView). Comparative analysis of complete genomes of *L. santarosai* serovar Shermani and genomes of pathogenic *Leptospira*s against the *L. santarosai* strain was performed by the Mallow alignment system and Blastp analysis. Protein functional domains were identified by searching the Pfam database.

Using the RAST programhttp://rast.nmpdr.org/, for the annotation of genomes, the functional annotation of the sequences reported for *L. santarosai* was sought, in addition, through this same program, the classification of said functional annotations in cellular subsystems was sought, where emphasis will be placed on the subsystem of factors of virulence.

### Search for polymorphisms between *Leptospira* species

A first part of the search for important genes that may be related to pathogenicity is the identification of polymorphisms between the contigs of the downloaded species. For this, the MUMmer package (http://mummer.sourceforge.net/) was implemented, in which there are pre-compiled programs of free access, and which are suitable for the alignment of genomes using direct information from the nucleotides, or translating the chains into proteins because there may be more conservation at the protein level than at the nucleotide content. For this case, the FASTA sequences of the contigs were used, and a general search for polymorphisms was carried out among all *Leptospira* species, following the following steps: Alignment between the contigs through the nucmer program, filtering the results to avoid repetitive regions, with the filter individual mutations were detected in order to map them to a list of proteins, taking into account the gene annotations obtained from GenBank.

The polymorphisms detected in certain regions are not necessarily linked to an event causing pathogenicity. Therefore, the polymorphisms were subsequently mapped with the list of orthologous proteins to infer at some level whether a mutation can be related to the expression or not of a protein in a pathogenic species compared to a non-pathogenic one.

### Analysis of orthologous proteins in pathogens that report polymorphisms with non-pathogenic species

Based on the prediction of orthologous proteins between different species, as well as the list of possible polymorphisms, a cross analysis was carried out to detect a list of genes that report orthologs only in pathogenic species including the Colombian strain *L. santarosai*, but which are not reported in species of intermediate pathogenicity or saprophytes. The purpose is to detect the presence of polymorphisms in the region that codes for the protein that is reported in pathogenic species, but not in saprophytic ones.

The proteins associated with these polymorphisms were mapped in the UniProtKB database19 and annotated information from the GeneOntology consortium was associated with each of the records in its sub-cellular localization component20.

### Search for orthologs between *Leptospira* species

The sequences in FASTA format of the three groups created from their association with pathogenicity were filtered following protocols established by the OrthoMCL5 program. The genes were identified and mapped through the GI code of the NCBI database. The results were grouped based on the presence of unique genes per pathogenicity category or by the presence of genes exclusively in certain categories and absence in the rest. The analysis allowed us to identify genes potentially associated with the pathogenicity of the bacteria. In addition, the program allowed identifying groups of orthologs in several categories: genes that reported at least one ortholog with another species, pathogen genes that did not have orthologs in non-pathogenic groups, genes that did not have orthologs in non-pathogenic groups and reported orthologs in all pathogenic and intermediate species, and total unique genes among all species.

For the group of genes from pathogens that did not have orthologs in non-pathogenic groups, that is, genes that were only found in pathogens, the cellular location of each of them will also be searched.For this purpose the proteins were mapped to the UniProt database (www.uniprot.org) and each of the records was searched for annotated information from the GeneOntology in its sub-cellular location component.

### Search for orthologs and unique genes between pathogenic and intermediate species

For this purpose, the OrthoMCL program was run with the *Leptospira* strains and species that had their proteins annotated.

### Analysis of the pangenome of the Colombian strain *L. santarosai* and different pathogenic, intermediate and saprophytic *Leptospira* strains and the pangenome of common genes between *L. santarosai* strains

The pan-genome is defined as a basic and common set of genes that are shared by all the genomes under study plus a variable set of genes shared by a subset of genomes, and specific or new genes of some strain22.

Taking into account the *L. santarosai* strains that are currently sequenced in the form of contigs, a search of the pangenome and core genome of the studied species was carried out. For this, the PanOCT program was implemented (http://sourceforge.net/p/panoct/home/Home/), which is a Perl script that not only looks for strategies to detect orthologs by conventional methods, but also takes into account the specific position in the genome of the coding sequences and the conservation of neighboring genes. To successfully run the program, it was necessary to create different files, taking into account protein information, such as the annotation files of the coding sequences (CDS) according to the Genbank format initially downloaded for each of the strains. The specific files that had to be created are the following: Multiple alignment between all strains through the blastp package of the NCBI BLAST+ toolkit (http://blast.ncbi.nlm.nih.gov/Blast.cgi), identifiers of each of the strains subjected to the algorithm, attribute file in which the contig is specified, the position in the genome, the ID and the function of the protein belonging to each of the strains analyzed and finally the protein sequences in FASTA format.

### Search for orthologs between the Colombian strain *L. santarosai* and the Taiwanese strain *L. santarosai* serovar Shermani str. LT 821

As an additional analysis of the above procedure to compare the number of orthologous genes between the Colombian strain *L. santarosai* and the Taiwanese strain *L. santarosai* serovar Shermani str. LT 821 (only *L. santarosai* strain assembled at the chromosome level)18 the BLAST algorithm was used to perform paired alignments between the previously downloaded sequences, in the case of *L. santarosai* serovar Shermani str. LT 821, proteins from both chromosomes were merged into a single file. Having the input FASTA files, the Inparanoid version 4.1.21 program was used to optimize the ortholog search protocol with the results obtained by BLAST.

Analysis of additional proteins of the Colombian strain *L. santarosai* based on comparison with genes reported in the literature of pathogenic species Taking into account a list of proteins associated with pathogenicity and reported in the literature mainly in *L. interrogans* strain Lai 56601 and *L. interrogans* strain Fiocruz L1-1130, a paired blastp was performed between the annotated proteins of *Leptospira santarosai* serogroup Autumnalis serovar Alice and the proteins of the reference species mentioned above. Genes with adequate percentages of identity to infer homology were searched in the UniProtKB database and information on their molecular function was searched to subsequently describe the pathogenic role of said bacterial protein in the disease.

## Results

### Sequencing, assembly and annotation

#### Nucleotide sequence accession number

The genome sequence of the strain *Leptospira santarosai* serogroup Autumnalis serovar Alice has been deposited in DDBJ/EMBL/GenBank under the accession number AKWS00000000.2.

#### Circular genome of the two chromosomes of the Colombian strain *L. santarosai*

Using the assembled chromosomes of *L. santarosai* serovar Shermani LT821 as a reference, the assembly at the scaffold level of the Colombian strain was constructed using circular maps generated by the CGview Comparison Tool program (Figure 1).

Chromosome 1: Chromosome 2:

**Figure 1.**
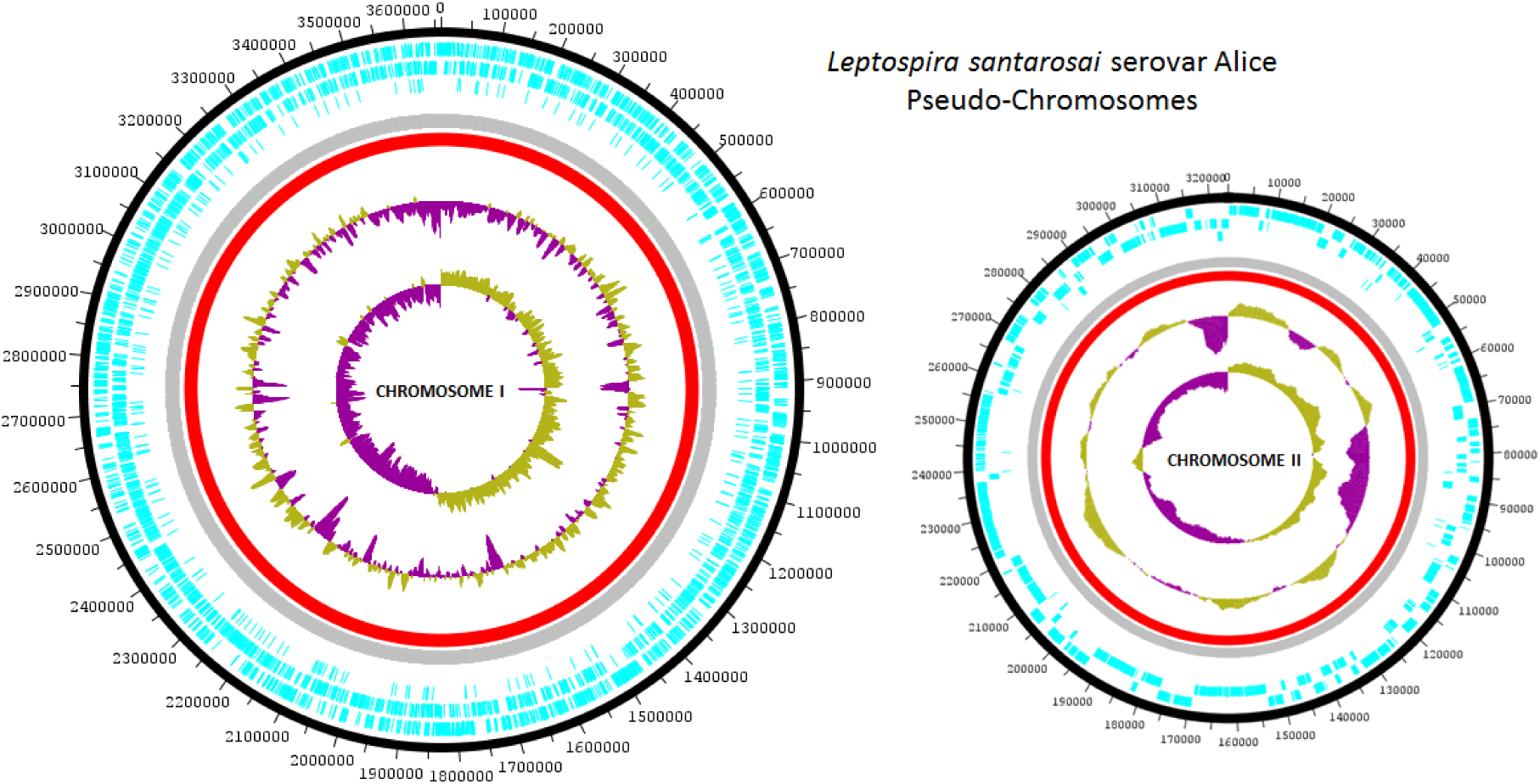
Circular maps of the chromosomes of the Colombian strain *L. santarosai*.

Using as reference the assembled chromosomes of *L. santarosai* serovar Shermani str. LT 821, the assembly at the scaffold level of the major (chromosome 1) and minor (chromosome 2) chromosomes of the Colombian strain *L. santarosai* is shown using circular maps generated by the CGview Comparison Tool program. The description of the circular map inside outward is the following, the innermost part is the inner scale where it is shown in Kilo base pairs (Kbp) indicating the size of the sequence. Then in purple the negative bias of the guanine and cytosine (GC) content, in green the positive bias of the GC content, in black the percentage of GC, in pink the comparisons made in Blast between the genome of the Colombian strain of interest of *L. santarosai* and the strain used as a comparison reference *L. santarosai* serovar Shermani str. LT 821 and finally in dark pink the open reading frames (ORF) are shown.

### Genome characteristics of the Colombian strain *L. santarosai*

The genome of *L. santarosai* Colombian strain consists of two circular chromosomes, the major chromosome and the minor chromosome, which add up to a total of 4.1 Mb. Circular representations of both chromosomes are shown in Figure 1. The two chromosomes of *L.* . *santarosai* have a G+C percentage of 41.6% containing 4540 coding sequences (CDS) of which 2145 (47.2%) have known function and the remaining 2395 (52.8%) have no described annotation. functional, its genome has 3918 genes of which 317 genes are specific to the Colombian strain *L. santarosai*, it has 4 genes for ribosomal RNA (rRNA) distributed 1 for the 23s subunit, 1 for the 16s subunit and 2 for the 5s subunit and 37 transfer RNA genes (Table 1).

**Table 1.**
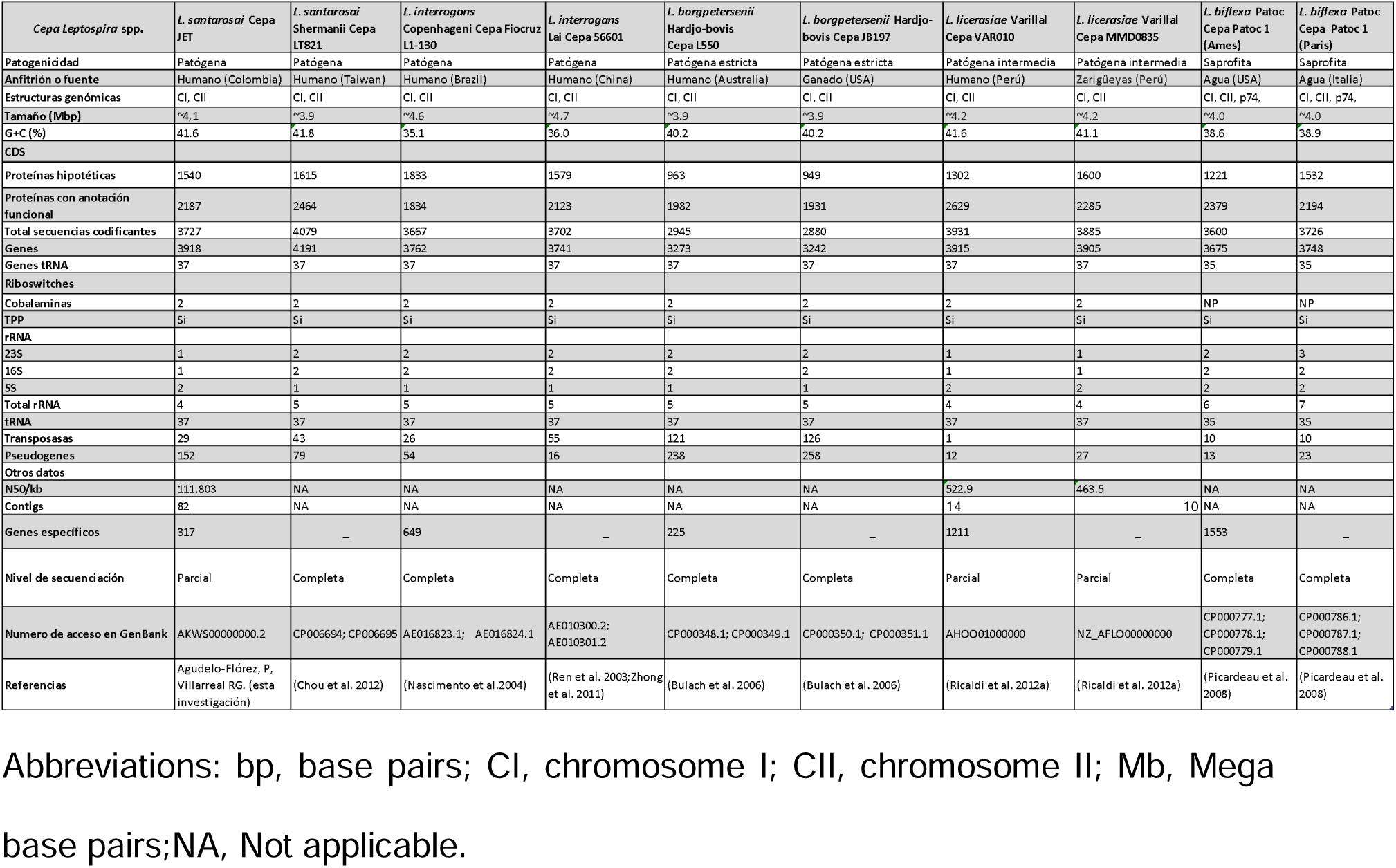
Genome characteristics of the Colombian strain compared to the reference strains *Leptospira* spp.

Using the RAST program and among the proteins for which an ortholog was found and a function could also be assigned, 85 proteins that are membrane components are reported, of which 17 are outer membrane proteins (Annex 1).

### Comparative genome analysis

The descriptive characteristics of the genomes of the strains compared with the Colombian strain of *L. santarosai* and the other reference strains in each group of pathogenic, intermediate pathogenicity and saprophytic *Leptospira* are found in Table 1.Where included: Pathogenicity, host or source, genomic structures, size (Mbp), G+C content (%), CDS coding sequences, hypothetical proteins, proteins with functional annotation, total coding sequences, genes, tRNA genes, riboswitches, cobalamins, TPP, rRNA, 23S, 16S, 5S, total rRNA, tRNA, transposases, pseudogenes among other characteristics. For the comparative analyzes all strains reported in Genbank were taken into account, which included pathogenic strains, intermediate pathogenicity strains, and saprophytic strains (Table 1).

### Annotation and subsystems of the Colombian strain *L. santarosai*

The functional annotation showed that the Colombian strain presents 4216 proteins of which 1831 (43.4%) have an assigned function and 2385 (56.5%) are hypothetical (Annex 1). Of the total proteins that were categorized into subsystems, 203 gene elements were found involved in bacterial metabolism, 197 in the production of amino acids, 152 in the formation of carbohydrates, 141 cofactors, 116 involved in the formation of cell wall, 103 in motility and chemotaxis, 99 involved in RNA metabolism and 96 involved in the formation of fatty acids and bacterial membrane lipids (Figure 2).

**Figure 2.**
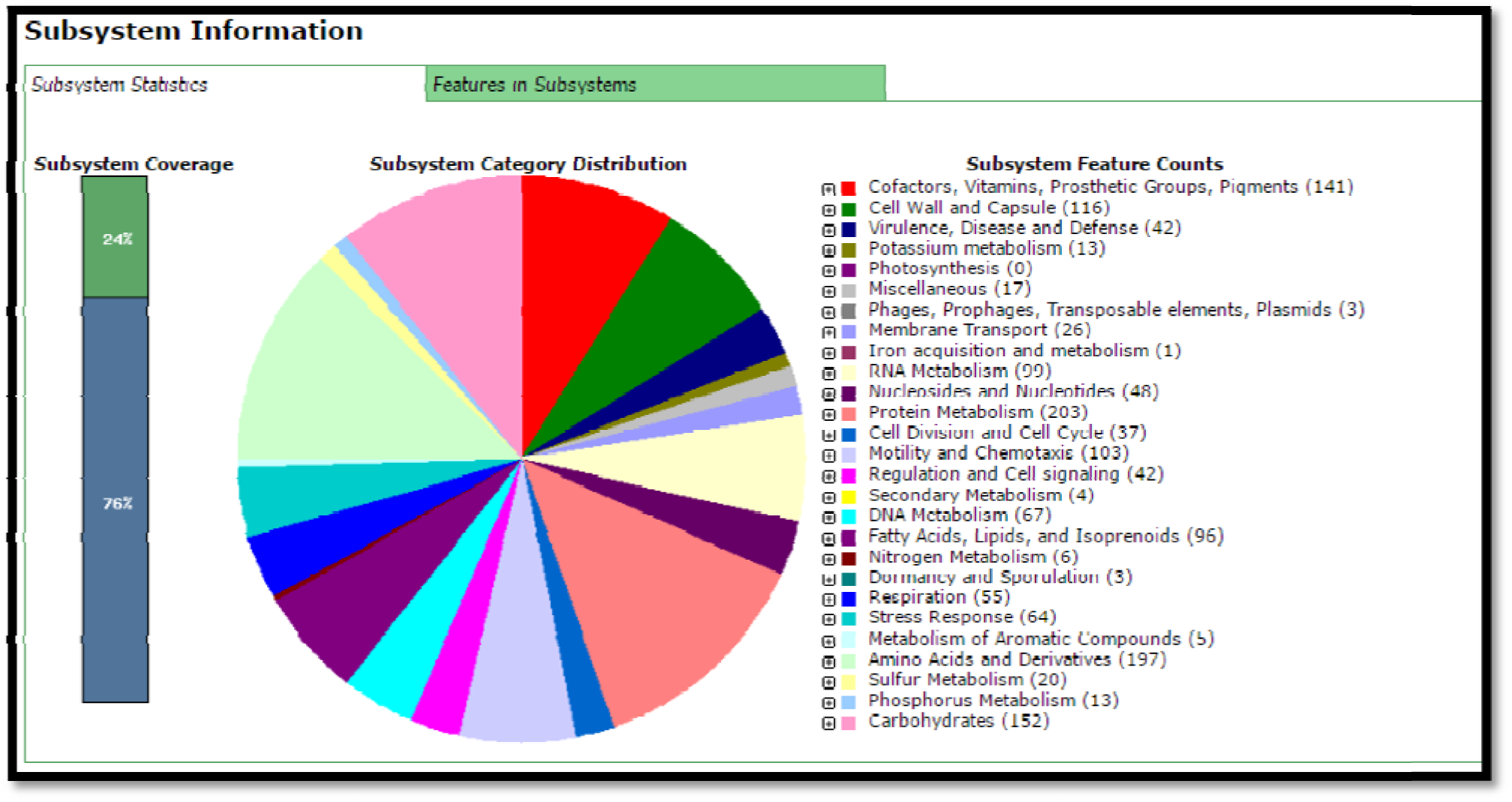
Subsystem information of the proteins annotated in the Colombian strain *L. santarosai*.

### Analysis of the pangenome of the Colombian strain *Leptospira santarosai* serogroup Autumnalis serovar Alice and different reference strains of pathogenic, intermediate and saprophytic *Leptospira*

Based on the genomes of the three analyzed subgroups of pathogenic, intermediate and saprophytic *Leptospira*, it was determined that the size of the core of the pangenome contained 1938 genes as these were the genes that were shared between all species, each subgroup in turn presents unique genes, and Shared genes are also evident between each of the subgroups.

Taking into account only the common genes between the strains of the different three groups, it is shown that the Colombian strain shares 2770 (75.8% similarity) orthologous proteins with the pathogenic strain *L.* interrogans, 2501 (68.5% similarity) with the intermediate strain *L.* licerasiae and 2106 (57.6% similarity) with the saprophytic strain *L.* biflexa. By demonstrating that the Colombian strain had a higher percentage of average identity with the pathogenic strains, this finding corroborates that the Colombian strain is more closely related to the pathogenic strains than to the saprophytic strains.It is also evident that there is a percentage of genes that did not present orthology with other genes; the majority of these genes encode hypothetical proteins. The distribution of clusters representing gene families among the three clusters is depicted in a Venn diagram in Figure 3.

**Figure 3.**
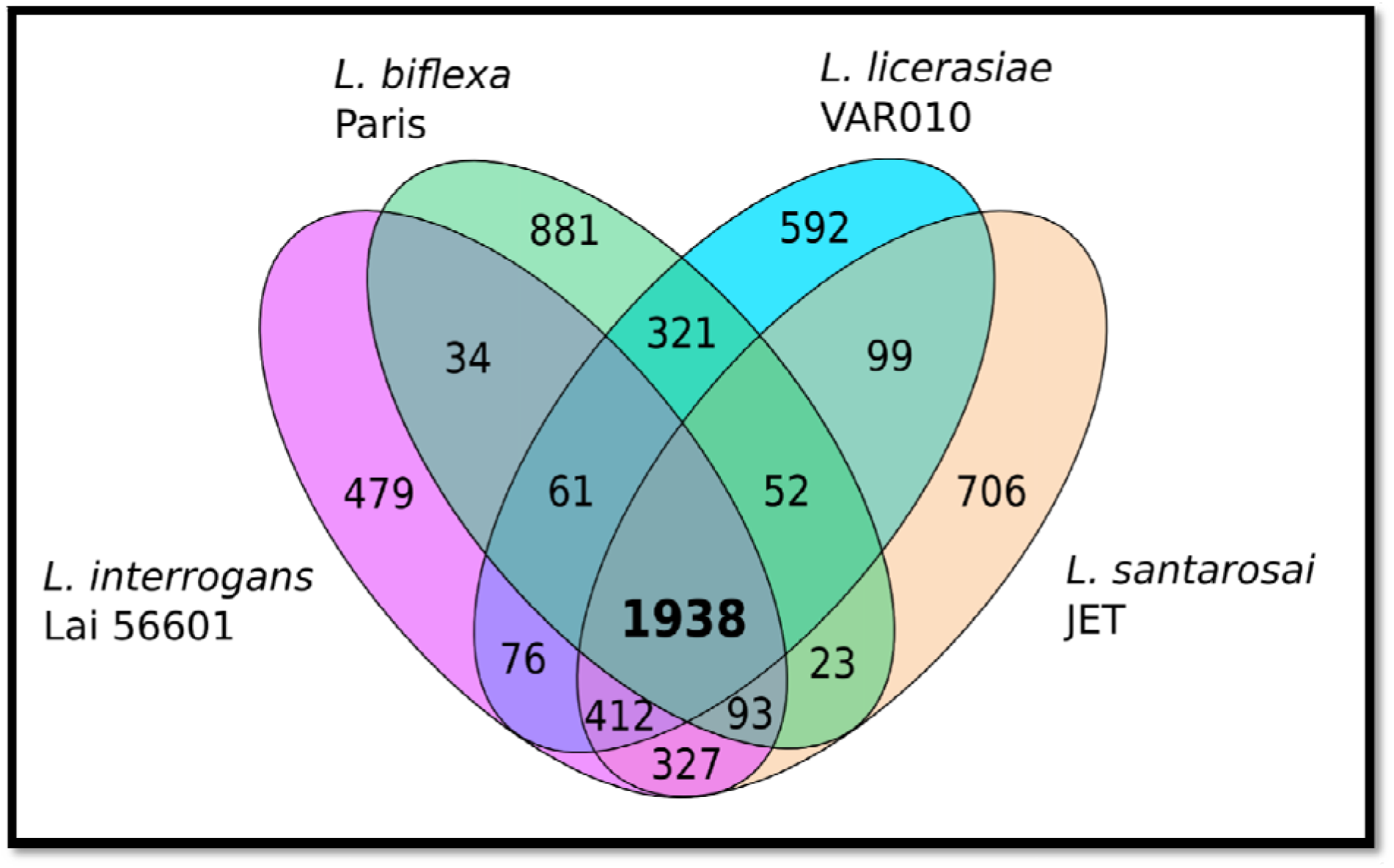
**venn diagram of**analysis of the pangenome distribution of *Leptospira santarosai* serogroup Autumnalis serovar Alice with strains from different pathogenic, intermediate and saprophytic *Leptospira* groups.

### Common genes in all pathogenic and intermediate *Leptospira* species

By including only genes found in pathogenic and intermediate strains and not including genes found in saprophytic strains, a total of 1738 common genes were identified between all intermediate and pathogenic species. Of these genes, 684 hypothetical or uncharacterized proteins were reported. These unique genes corresponded to genes that encode cytoplasmic and intracellular proteins and membrane components (Appendix 2).

### Search for orthologs between the Colombian strain *Leptospira santarosai* serogroup Autumnalis serovar Alice and the *L. santarosai* strains with sequenced genome

Because the interest of the project lies in *L. santarosai*, information related to various strains of this species was also downloaded, among which were the Colombian strain *Leptospira santarosai* serogroup Autumnalis serovar Alice and another strain reported in Colombia characterized as *L. santarosai* AIM so it is also of great interest, all of these strains were annotated at the contig level (Table 2). In this case, substantial differences are observed in the number of contigs, and in the size in Mb of the genomes. The two Colombian *L. santarosai* strains analyzed in this study are intermediate, with N50 values ranging between 111,000 and 121,000 base pairs. Despite not being the best annotation, the combination of Illumina and 454 technology allows covering a significant amount of genomic information in order to perform the following analyses.

**Table 2.**
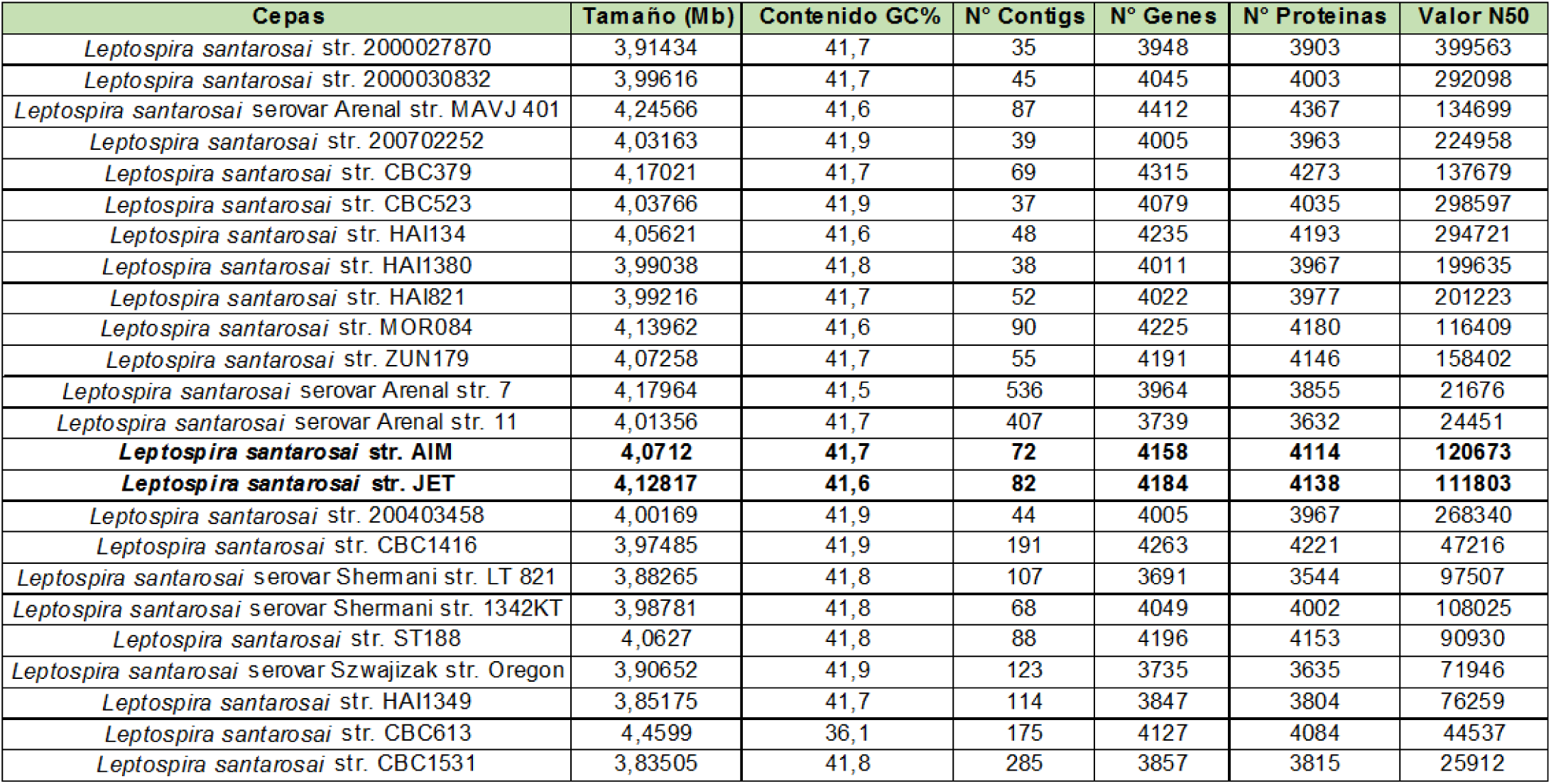
List of *L. santarosai* strains selected for analyses. *The two Colombian strains of *L. santarosai* are highlighted in black.

| Cepas | Tamaño (Mb) | Contenido GC% | N° Contigs | N° Genes | N° Proteínas | Valor N50 |
| --- | --- | --- | --- | --- | --- | --- |
| *Leptospira santarosai* str. 2000027870 | 3,91434 | 41,7 | 35 | 3948 | 3903 | 399563 |
| *Leptospira santarosai* str. 2000030832 | 3,99616 | 41,7 | 45 | 4045 | 4003 | 292098 |
| *Leptospira santarosai* serovar Arenal str. MAVJ 401 | 4,24566 | 41,6 | 87 | 4412 | 4367 | 134699 |
| *Leptospira santarosai* str. 200702252 | 4,03163 | 41,9 | 39 | 4005 | 3963 | 224958 |
| *Leptospira santarosai* str. CBC379 | 4,17021 | 41,7 | 69 | 4315 | 4273 | 137679 |
| *Leptospira santarosai* str. CBC523 | 4,03766 | 41,9 | 37 | 4079 | 4035 | 298597 |
| *Leptospira santarosai* str. HAI134 | 4,05621 | 41,6 | 48 | 4235 | 4193 | 294721 |
| *Leptospira santarosai* str. HAI1380 | 3,99038 | 41,8 | 38 | 4011 | 3967 | 199635 |
| *Leptospira santarosai* str. HAI821 | 3,99216 | 41,7 | 52 | 4022 | 3977 | 201223 |
| *Leptospira santarosai* str. MOR084 | 4,13962 | 41,6 | 90 | 4225 | 4180 | 116409 |
| *Leptospira santarosai* str. ZUN179 | 4,07258 | 41,7 | 55 | 4191 | 4146 | 158402 |
| *Leptospira santarosai* serovar Arenal str. 7 | 4,17964 | 41,5 | 536 | 3964 | 3855 | 21676 |
| *Leptospira santarosai* serovar Arenal str. 11 | 4,01356 | 41,7 | 407 | 3739 | 3632 | 24451 |
| ***Leptospira santarosai* str. AIM** | **4,0712** | **41,7** | **72** | **4158** | **4114** | **120673** |
| ***Leptospira santarosai* str. JET** | **4,12817** | **41,6** | **82** | **4184** | **4138** | **111803** |
| *Leptospira santarosai* str. 200403458 | 4,00169 | 41,9 | 44 | 4005 | 3967 | 268340 |
| *Leptospira santarosai* str. CBC1416 | 3,97485 | 41,9 | 191 | 4263 | 4221 | 47216 |
| *Leptospira santarosai* serovar Shermani str. LT 821 | 3,88265 | 41,8 | 107 | 3691 | 3544 | 97507 |
| *Leptospira santarosai* serovar Shermani str. 1342KT | 3,98781 | 41,8 | 68 | 4049 | 4002 | 108025 |
| *Leptospira santarosai* str. ST188 | 4,0627 | 41,8 | 88 | 4196 | 4153 | 90930 |
| *Leptospira santarosai* serovar Szwajizak str. Oregon | 3,90652 | 41,9 | 123 | 3735 | 3635 | 71946 |
| *Leptospira santarosai* str. HAI1349 | 3,85175 | 41,7 | 114 | 3847 | 3804 | 76259 |
| *Leptospira santarosai* str. CBC613 | 4,4599 | 36,1 | 175 | 4127 | 4084 | 44537 |
| *Leptospira santarosai* str. CBC1531 | 3,83505 | 41,8 | 285 | 3857 | 3815 | 25912 |

The core-genome among all *L. santarosai* strains is composed of 1747 proteins, which are common to all *L. santarosai* strains analyzed. In the case of the Colombian strain *Leptospira santarosai* serogroup Autumnalis serovar Alice, of the total of 4138 proteins, 141 are exclusive to its genome and 13 are shared with the *L. santarosai* AIM strain, which reports 158 exclusive proteins. This indicates that approximately each *L. santarosai* strain may contain approximately 3.5% exclusive genes, which may be related to its infectivity or resistance mechanisms. A general distribution of orthologs between the strains of interest and the others is shown in Figure 4.

**Figure 4.**
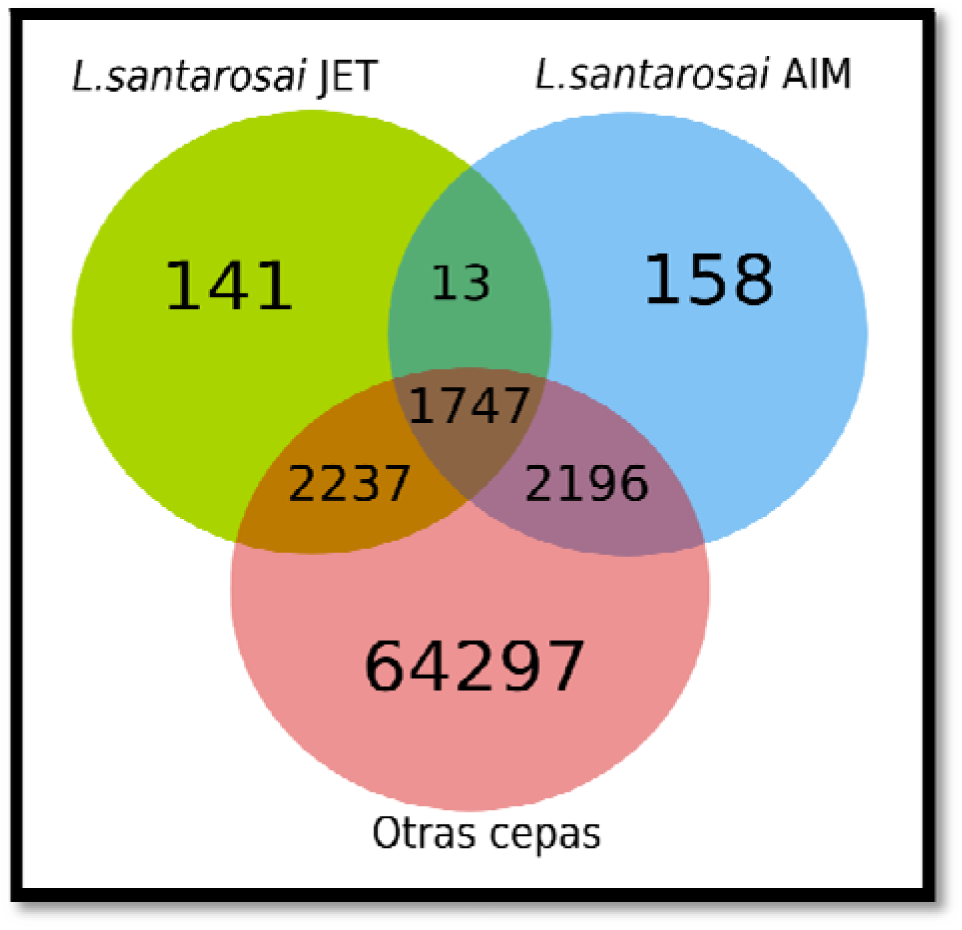
Venn diagram with the distribution of common or unique genes according to pangenomic analysis among the different strains of *L. santarosai*.

### Search for orthologs between the Colombian strain *Leptospira santarosai* serogroup Autumnalis serovar Alice and the Taiwanese strain *L. santarosai* serovar Shermani str. LT 821

The strain *L. santarosai* shermani str. LT821 is the only strain of the *L. santarosai* species for which the genome has been completely sequenced18, making it important as a reference for the comparative analysis of this study. There are several differences ranging from genetic characteristics to variability in the similarity of orthologous proteins between the two *Leptospira* species. The Colombian strain has a size of 4.1 Mpb and the Taiwanese strain has a size of 3.9 Mpb, the Colombian strain has a total of 3918 genes and the Taiwanese strain 4191 genes, the Colombian strain has a total of 3727 proteins coding and the Taiwanese strain 4079, the Colombian strain has 2187 (58.6%) proteins with functional annotation and the Taiwanese strain 2464 (60.4%), the Colombian strain has 1540 (41.3%) mortgage proteins while that the Taiwanese strain presents 1615 (39.5%) (Table 1).

Regarding the number of orthologous genes that the two strains present, it can be see that 3262 orthologous genes are shared between them, which represents 78% similarity, and that the Colombian strain has 876 different genes, while the Colombian strainTaiwaneseIt presents a total of 818 different genes (Figure 5).

**Figure 5.**
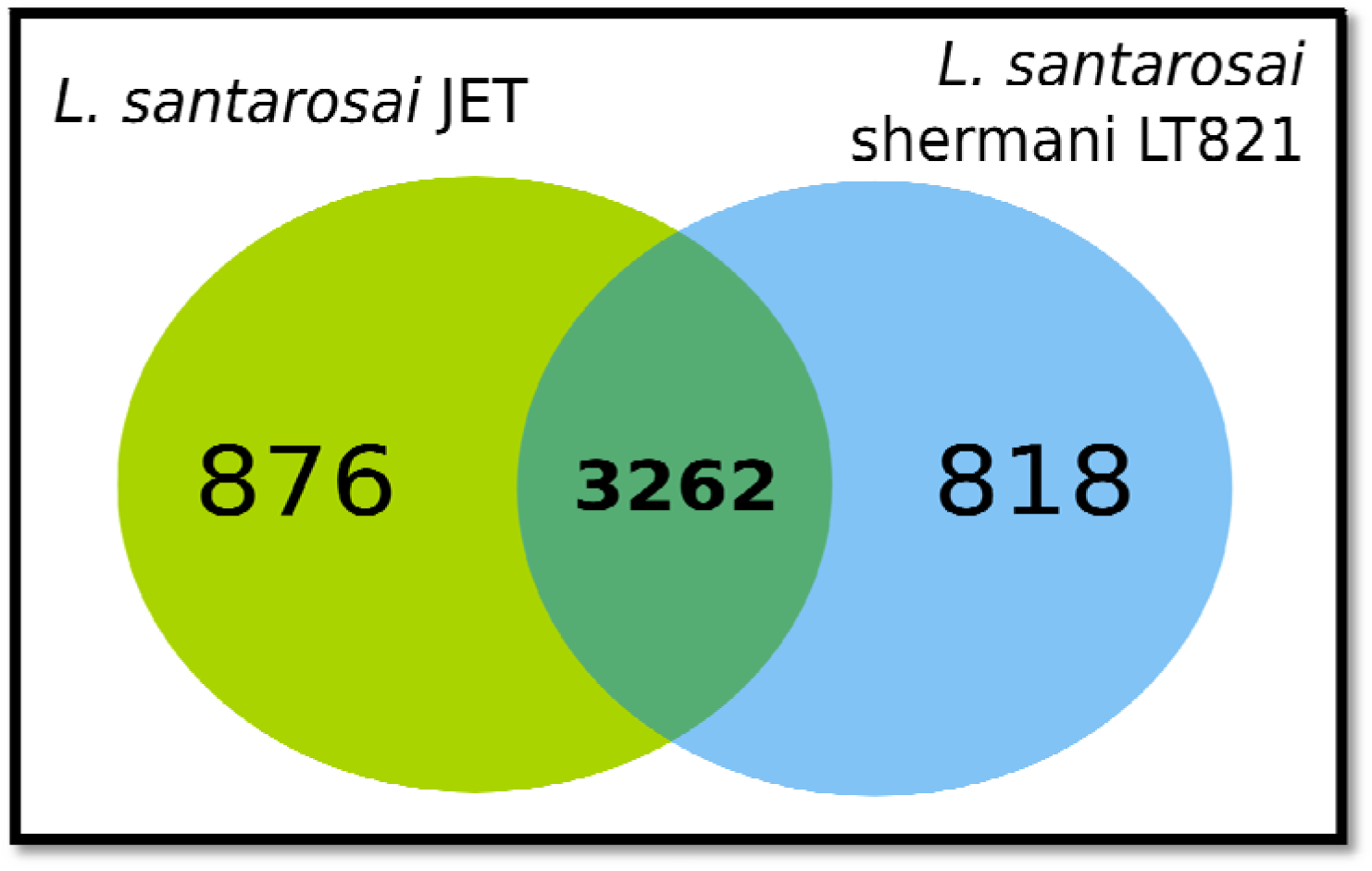
Venn diagram of orthologs between the Colombian strain *Leptospira santarosai* serogroup Autumnalis serovar Alice and the Taiwanese strain *L. santarosai* serovar Shermani str. LT 821.

### Search for orthologs between *Leptospira* species

From the general analysis, the following statistics were obtained for the Colombian strain *L. santarosai*: 317 unique genes, 3650 genes reported at least one ortholog with another species, 959 genes that had no orthologs in non-pathogenic groups and 37 genes reported orthologs in all species pathogenic and had no orthologs in intermediate and saprophytic *Leptospira* groups.

Taking into account the previous groups of genes detected, the one that draws special attention is that group in which orthologous genes are found only in pathogenic *Leptospira* species. This list yielded a total of 37 proteins, of which a group were hypothetical (uncharacterized) proteins, and 17 were proteins with reported functions, which may play an important role in the pathogenicity of the strain (Table 3). This group of proteins was assigned its identification code, description, function, possible role in virulence depending on its function, subcellular location and references from previous works on these proteins where possible functions were deduced through sequencing and experimental confirmations of some of them. Several proteins were poorly studied and no information was reported (NR).These results highlight the presence of several copies of the sphingomyelin phosphodiesterase protein, which may play an important role in the molecular mechanisms used by the bacteria to infect. The functional domains or families detected and other characteristics of these proteins are shown in Table 3.

**Table 3.** List of proteins with known function from *L. santarosai* that report orthologs with all pathogenic species and do not report orthologs with intermediate or saprophytic species.

| GI proteína | Descripción/Función | Ubicación | Posible rol en virulencia | Referencia |
| --- | --- | --- | --- | --- |
| 410803357 | PF07603 family protein (Xylanase) Degradación de carbohidratos | N.R | Adquisición de nutrientes | Hartskeerl R.A 1et al. 2011 |
| 410804148 | Trypsin like peptidase domain protein Rompe los enlaces peptídicos de las proteínas mediante hidrólisis para formar péptidos de menor tamaño | Transmembrana | Daño epitelial | Hospenthal D et al. 2011 |
| 410803664 | Putative sphingomyelin phosphodiesterase Actividades hemolíticas | Región Extracelular | Invasión de tejidos, daño endotelial, evasión inmune y adquisición de nutrientes | Narayanavari et al. 2012. Nascimento et al. 2004 |
| 410803657 | Putative lysophospholipase | N.R | Adquisición de nutrientes/ Daño epitelial | Yanagihara Y et al. 1984 Nascimento et al. 2004 |
| 410808148 | Sigma factor regulatory protein, FecR/PupR family FecR participa en la regulación del transporte dicitrato hierro. Internalización del hierro | Transmembrana | Adquisición de nutrientes | Weiz D e al. 1998 |
| 410803096 | Sigma-70 region Z Factor sigma necesario para transcripción del flagelo | N.R | Diseminación | Lin M et al. 2004. |
| 410802828 | putative membrane protein | Transmembrana |  |  |
| 410802399 | Dienelactone hydrolase family protein Hidrolasa | N.R | Adquisición de nutrientes | He P et al.2013 |
| 410802315 | sphingomyelin phosphodiesterase | Región Extracelular | Adquisición de nutrientes | Narayanavari et al. 2012. Nascimento et al. 2004 |
| 410802207 | tRNA pseudouridine synthase D (enzimas responsables de la modificación postranscripcional más abundante de los ARN celulares) | N.R | Metabolismo | Hamma T et al. 2006 |
| 410802098 | FG-GAP repeat protein (lipoproteína involucrada a la adhesión celular) | Complejo Integrina | Entrada en la célula | Picardeau M et al. 2012 |
| 410801350 | STAS domain protein (Sulphate Transporter and Anti-Sigma factor antagonist, domain of anti-anti-sigma factors) | N.R | Adaptación | Eshghi A et al. 2014 |
| 410800916 | Glucose-methanol-choline-GMC oxidoreductase (Actividad oxidoreductasa, que actúan sobre el grupo CH-OH de los donantes, otros aceptadores) | Transmembrana | Evasión del sistema inmune | Geo J et al. 2014 Prêtre G et al. 2011 |
| 410800890 | Methyltransferase domain protein<br>transducción de señal quimiotáctica en bacterias | N.R | Motilidad y expresión de proteínas | Sun YJ et al. 2005<br>Batra M al. 2015 |
| 410800886 | Bacteriocin-protection protein,<br>Ydel/OmpD-associated family<br>Confiere protección contra AMP<br>catiónicos (péptidos antimicrobianos) o<br>bacteriocinas en conjunción con la<br>porina en general.<br>Mecanismos de autoprotección | N.R | Supervivencia | Mathur H et al. 2015 |
| 410800850 | AAA ATPase domain protein<br>Mecanismo Inflamatorio | Intracelular | Daño epitelial | Lacroix-Lamande S et al. 2012 |
| 410800823 | Sphingomyelin phosphodiesterase | Región Extracelular | Adquisición de nutrientes | Narayanavari et al. 2012.<br>Nascimento et al. 2004 |

## Discussion

In the present work, the genome of *Leptospira santarosai* serogroup Autumnalis serovar Alice was sequenced, circular maps of the major and minor chromosomes of this bacterium were generated and comparisons were made with other pathogenic, intermediate and saprophytic *Leptospira* genomes to search for orthologs. used the analysis of the sequences, their domains and their structures to elucidate the functions of the proteins and description of hypothetical proteins, finding notable similarities with respect to the pathogenic strains, which was used to find virulence factors19 involved in the pathophysiology of this Colombian strain.Also, thisPangenome also reflected the high genomic variability between *Leptospira* species and intraspecies as in the case of *L. santarosai*. This genomic diversity can be attributed to the gain and loss of genes, reporting greater genetic losses in pathogenic strains compared to saprophytic strains, which deduces that each species makes high changes in its genetic composition to adapt to specific conditions of variable environments. According to this study, it is inferred that the Colombian strain has unique genes acquired through horizontal transfer. These genes may in turn allow it to express unique virulence factors, but for this hypothesis it is necessary to carry out subsequent experimental validations. When analyzing the statistics of the proteins expressed by the Colombian strain in cellular subsystems, it was found that the majority of its genome expresses genes involved in the generation of proteins necessary for cellular metabolism, amino acids, carbohydrates, enzymatic cofactors, cell wall, motility and chemotaxis., fatty acids and lipids. It should be noted that many of these genes encode components of the outer membrane or compounds associated with the membrane, and because the membrane components are the first thing exposed by the bacteria and are the main ones to come into contact with the host cells, this would indicate that the Colombian strain of *L. santarosai* uses a significant amount of its genetic material in the interaction with the host cell.

Comparative genomics approaches began to provide insight into the large set of genes that allow pathogenic *Leptospira* strains to adhere, invade, persist, evade the immune system, colonize and cause disease in mammalian reservoirs, as well as accidental hosts.

Through the identification of new gene families, differences and similarities can be established between the different groups of *Leptospira* that can be important, especially in pathogenic groups, by being able to elucidate the mechanisms of pathogenicity of the bacteria. In the present work, it was demonstrated that pathogenic *Leptospira* species contain unique genes that are not found in saprophytic *Leptospira*. In this way, 37 genes exclusive to pathogenic species were found, of which 17 could be assigned a function.

With these virulence factors found in the Colombian strain *L. santarosai* serogroup Autumnalis serovar Alice and that are found alone and all pathogenic species of *Leptospira*, the description of thestages in *Leptospira* infection, since virulence factors are necessary in the pathogenesis of *Leptospira* initially for the entry and dissemination of the bacteria22,23, which enters through the mucosa and eroded skin, here the motility of the bacteria is necessary where Proteins such as Sigma- 70 region 2 intervene in the Colombian strain to increase its flagellar movement and be able to advance within the host.

Chemotaxis is another important property because it allows the bacteria to direct itself towards a high concentration gradient of nutrients essential for its survival22,23. Here, proteins such as Methyltransferase domain protein 2 were found that facilitate chemotaxis and allow it to direct itself towards a specific substrate and be able to acquire food sources. This protein is a chemoreceptor that helps increase the movement of the flagellum according to the increased concentration gradient of nutrients in the medium. These proteins may be essential for their survival in specific environments, including their ability to infect a host.

Once the bacteria enters, it needs to adhere to the extracellular matrix22,23 at this point different proteins have been found in the Colombian strain. These proteins interact with extracellular matrix proteins, the FG-GAP repeat protein, which is a lipoprotein involved in binding to fabric.

The bacteria then have to penetrate and degrade tissue.^22,23^, at this stage several proteins were found such as Trypsin like peptidase domain protein, Putative sphingomyelin phosphodiesterase, Putative lysophospholipase involved in these tissue degradation functions to enter the tissue to reach blood vessels.

Once the bacteria manages to reach the bloodstream, the bacteria must persist, acquiring nutrients and evading the immune response22,23. Proteins necessary to acquire nutrients such as iron from erythrocytes as well as lipids from cell membranes were found in the Colombian strain. These proteins were PF07603 family protein, Putative lysophospholipase, Sigma factor regulatory protein, FecR/PupR family, Dienelactone hydrolase family protein and Sphingomyelin phosphodiesterase. . To persist, the bacteria also have to fulfill metabolic functions22,23; in this strain, proteins involved in metabolic processes such astRNA pseudouridine synthase D, STAS domain proteinandMethyltransferase domain protein.The Bacteriocin-protection protein involved in theSelf-protection which gives protection against cationic AMPs (antimicrobial peptides) or bacteriocins.Another characteristic necessary to persist is the evasion of the immune response22,23. Proteins were found in the Colombian strain that inhibit the complement cascade and prevent death by phagocytes by preventing oxidative stress. These proteins are Putative sphingomyelin phosphodiesterase and Glucose-methanol- choline-GMC oxidoreductase.

Finally, the bacteria spread to many organs such as kidney, lung, liver, muscle and even the central nervous system and aqueous humor to produce disease or even death of the host.^22,23^.

The above virulence factors found may be essential in the pathogenesis of the bacteria and may be useful in the creation of molecular targets for specific diagnosis, new treatments by using them as targets for future drugs and possible vaccines that are relevant to public health. humans and animals of veterinary importance. This work contributes to new knowledge about the evolution of a bacterial pathogen such as *Leptospira*, providing new molecular contributions to describe the pathogenesis of leptospirosis through its infective route in a host 24,25.

## Financing

This project was funded by Colciencias project 325649326207-678 122865740423.

## Acknowledgments

We thank Dr. Piedad Agudelo-Flórez for the methodological advice and the CES University for the loan of facilities for the execution of the project.

